# Smartphone-Operated Affordable PCR Thermal Cycler for the Detection of Antimicrobial Resistant Bacterial Genes

**DOI:** 10.1101/2022.05.27.493737

**Authors:** Sanam Pudasaini, Garima Thapa, Bishnu P. Marasini, Basant Giri

**Affiliations:** Center for Analytical Sciences, Kathmandu Institute of Applied Sciences, Kathmandu, Nepal; Department of Biotechnology, National College, Tribhuvan University, Naya Bazar, Kathmandu, Nepal

**Author notes:** **Correspondence:** Basant Giri, PO Box: 23002.

**Keywords:** Antibiotic resistance, *Escherichia coli*, Polymerase chain reaction (PCR), Arduino, resource limited setting, global public health threat, *bla-TEM*, *bla-CTXM*

## Abstract

**Objectives:** Antimicrobial resistance (AMR) is a global public health threat. Surveillance of AMR requires affordable, rapid, and user-friendly diagnostic method. Our aim was to develop a low-cost thermocycler to perform polymerase chain reaction (PCR).

**Methods:** We developed a smartphone-operated PCR thermal cycler using locally available recycled materials. The thermal cycler was used for the amplification for three bacterial genes - *bla-TEM* and *bla-CTXM and* 16s rRNA in human urine samples. The performance of custom-built thermal cycler was compared with the commercial one.

**Results:** The thermal cycler was portable (<1kg weight), required 12 V power supply, 25 µL of solution, and cost only USD50.0. Temperature and time conditions were instructed using a custom-built smartphone application. The ramping rate of was 0.23°C for heating and 0.43°C for cooling, set temperatures were within ± 0.5 °C of target showing a good thermal stability. The antibiotic sensitivity test of human urine samples showed they were highly resistance and multi-resistant. Nearly 46 % (n=54) *E. coli* isolates were positive in ESBL screening test. The custom-built thermocycler was able to accurately predict the presence of *bla-TEM* and *bla-CTXM* genes (n=6).

**Conclusions:** We developed and demonstrated a portable, low-cost, easy-to-use, and smartphone-operated PCR thermal cycler. Since it is portable, it can be used in remote location and field settings, including places without stable power supply. The use of the thermal cycler system can be extended, beyond the detection of AMR genes, e.g., in clinical diagnosis, genetics, forensic analysis, and environmental protection.

## 1. Introduction

Antimicrobial resistance (AMR) is one of the top 10 global public health threats [1]. According to the World Health Organization (WHO), AMR significantly impacts economy and is a threat to development. AMR occurs when microbes such as bacteria, virus, fungi, and parasites change over time and do not respond to antimicrobial drugs[1]. Resistant and multi-resistant bacteria have been detected in foods, farms, industries, slaughterhouses[2]. Surveillance of AMR to obtain a realistic threat measurement is one of the approaches to address this complex problem. There are several methods of identifying resistant bacteria and their respective resistance genes. The culture based traditional methods are simple and easy to carry out but require long time and do not work for non-cultivable microorganisms. Recently, molecular methods such as polymerase chain reaction (PCR) that are based on the amplification of target genes have been used in laboratory routine use for detecting AMR genes in multiple environments. [2]

Polymerase chain reaction (PCR) is a molecular technique used to amplify small segments of DNA [3]. After its introduction in 1985 [4], this technique has enabled scientific advancement in the field of medical science, forensic science, drug discovery, pathogen diagnosis, cell biology, biotechnology, and environmental protection [5,6]. PCR is usually performed using repeated thermocycling stages where different temperatures are used for different reaction stages such as denaturation, annealing, and extension in a single amplification cycle. During denaturation, two strands of DNA templates are separated. Similarly, during annealing step, primers bind to flanking region of DNA templates whereas during extension step, new DNA strands are formed[7]. To ensure proper amplification, PCR mix including polymerase enzymes, primers, targeted DNA templates and specific buffers needs to be optimized for a given application [8]. PCR plays a critical role in identifying gene responsible for developing antibiotic resistance in microorganisms.

Recently, many low-cost PCR systems have been developed for various applications. Additionally, many homemade control systems have been proposed that comprise multiple customized hardware to achieve the goal [10]. Hardware and software customizations not only increase the cost of the PCR equipment, but also have a technical barrier affecting popularization of the technology. Similarly, few microfluidic channels and chambers have been developed to serve as a platform for conducting PCR reactions[11–13]. Although faster reactions were reported, bulky external support systems such as pumps and tubing are required to inject the samples into the microfluidic device making the process complex in out-of-lab settings [14,15]. In addition, surface treatment is crucial in microfluidic system to prevent non-stick binding of samples and to prevent evaporation of samples[16,17]. Liu et al. developed a microfluidic device integrating numerous hydraulic valves and pneumatic pumps to deliver 2 µL of PCR mixture for around 400 independent reactors[18]. However, the implementation of valves and pumps is a complex fabrication process and can lead to expensive PCR operation [19]. Similarly, Kamalalayam et al. demonstrated the use of liquid marble that is coated with hydrophobic/oleophobic powder that acts as a reactor for PCR reaction[20]. Use of liquid marble eliminates the possibility of contamination as well as reduces the use of plastic waste. However, preparation of liquid marble is quite challenging, and expensive.

In last few decades, many commercial PCR systems have been introduced with excellent performance in amplifying DNA fragments of both animal and plant origin. The molecular methods for AMR detection can be executed in less time and can identify all resistomes in a sample, have high specificity and equally used for non-cultivable organisms.[2] But they are more expensive than traditional cultivation method and require specific equipment, technicians, and expertise limiting their wider use, especially in resource limited laboratories[9]. Even though the cost of molecular technology is decreasing significantly in the past decade, the cost of equipment and reagents is still high, especially for low- and middle-income countries (LMCs). The LMCs also lack trained personnel since educational institutions are not able to train their graduates on these technologies due to cost associated with it. Hence, at this point, the development of affordable PCR system is of great value in LMCs for both human and plant diseases surveillance and educational purposes. Availability of affordable, rapid, and easy to use PCR system for detecting AMR would address a critical need in surveillance of AMR globally.

Here, we developed a thermocycler at a very low cost (material cost <$50). The thermocycler used a proportional-integral-derivative (PID) controller to precisely control the temperature and was controlled and operated by a smartphone program using a bluetooth communication. After the development, we demonstrated the application of the system by amplifying three genes - rRNA encoding DNA (rDNA), *bla-TEM* and *bla-CTX-M* that produce extended spectrum beta lactamases (ESBL) in *E. coli*. A recent study has demonstrated a higher prevalence rate of ESBL producing *Escherichia coli* (*E. coli)* bacteria isolated from urine samples from a hospital in Nepal. We also compared amplification of genes using our custom developed low-cost thermocycler with a conventional PCR system.

## 2. Material and methods

### 2.2 Chemicals and reagents

The PCR solution (25 µL) contained 1 µL forward primer, 1 µL reverse primer, 10 µL DI water, 1 µL template DNA, and 12 µL NEB master mix (NEB, United Kingdom). Nuclease free water (GeNei, India) was used as a negative control.

### 2.2 Design and fabrication of thermocycler

A custom-built thermocycler was developed to run PCR thermal cycles. Figure 1A shows a schematic diagram of the custom-built thermocycler whereas actual fabricated equipment is presented in Figure 1B. A 35 × 40 × 30 mm^3^ aluminum block was used for holding PCR tubes. 40W cartridge heater (Core electronics) was embedded on an aluminum block and was used as a heating platform. The block was placed on the top of a 40 × 40 × 3.5 mm^3^ Peltier thermoelectric cooler (TEC-12706, AUS electronics). The Peltier device was used as a cooling platform. Thermal paste was applied between the aluminum block and Peltier device to enhance heat conduction. Entire assembly was mounted on the top of the heat sink to release heat from the hot side of the Peltier device. A computer CPU cooling fan was used to cool the heat sink. Thermocouple module was used to sense the temperature of the aluminum block. A PID algorithm was implemented on Arduino uno board.

**Figure 1:**
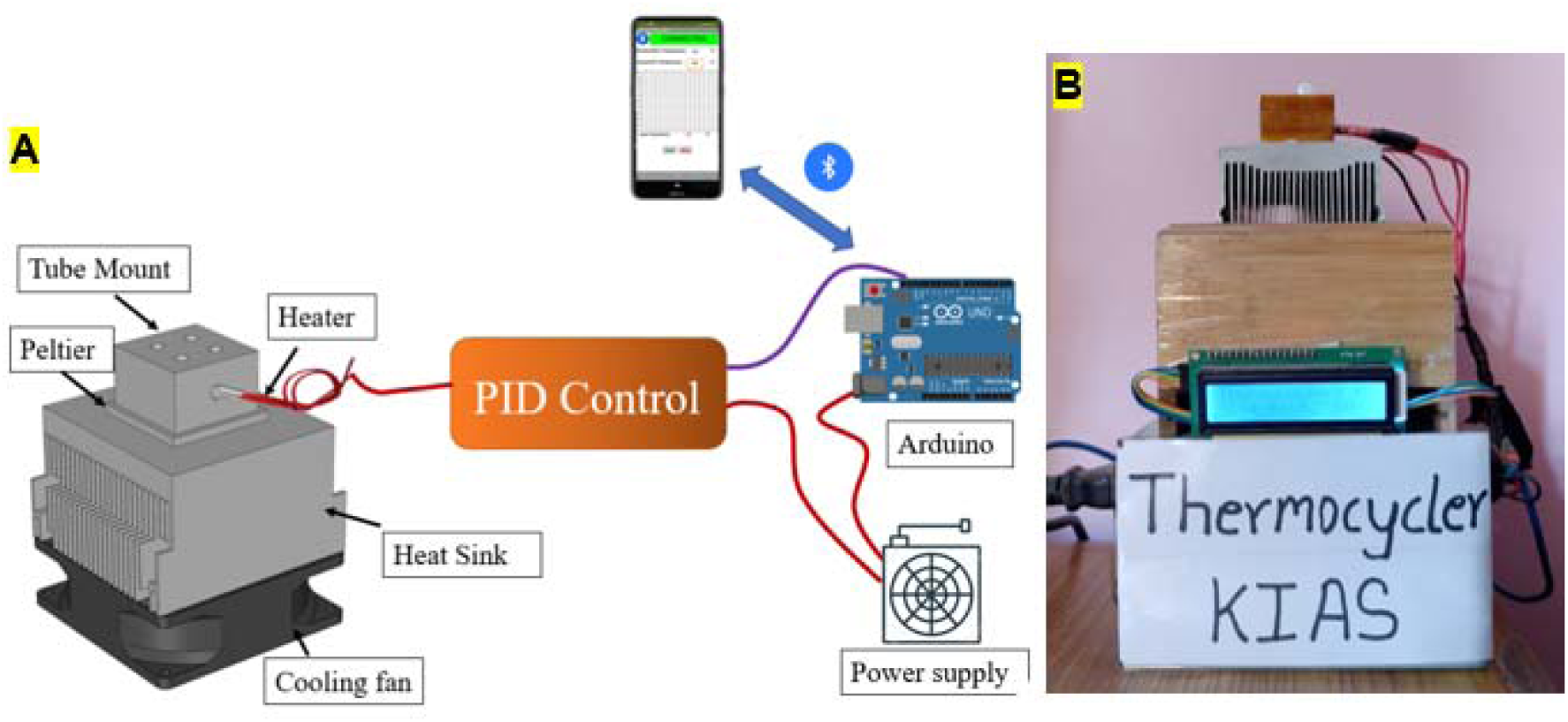
(A) Schematic diagram of low-cost PCR system. Smartphone sends temperature and time conditions to Arduino UNO via Bluetooth communication. Arduino then controls heating and cooling devices using the PID algorithm. (B) Photograph of the custom-built thermocycler assembly.

A smartphone application was developed using MIT app inventor[21] to provide temperature and process time via Bluetooth communication to Arduino uno. Meanwhile, the mobile application also received temperature data from the Arduino and plotted a graph of the temperature cycle. Figure 2A shows the smartphone screen of the custom developed application and detailed electrical diagram is presented in Figure 2B.

**Figure 2:**
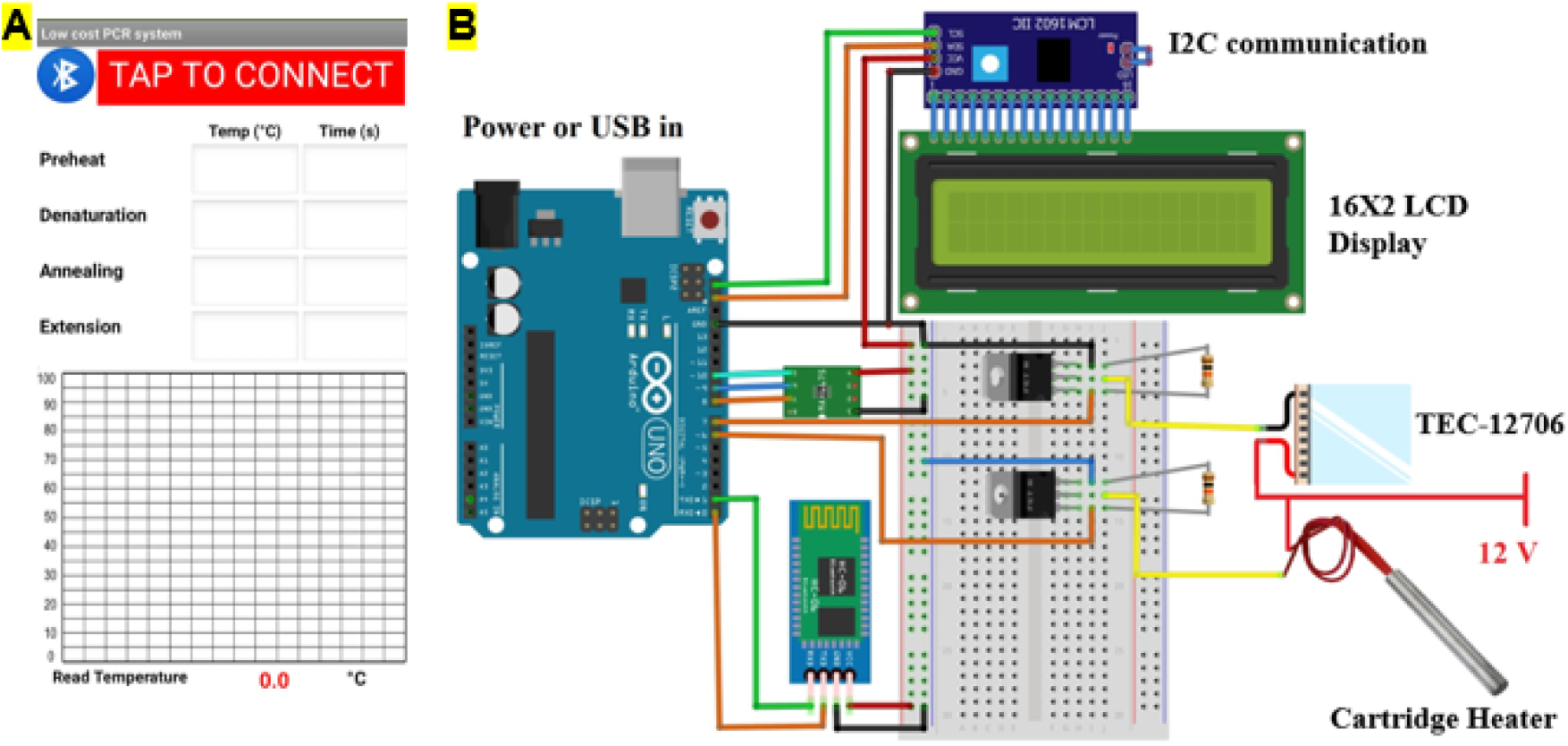
(A) Mobile screen presenting custom developed application for our thermocycler. The app sends temperature and time data to the Arduino via Bluetooth communication and receive real time temperature data from the thermocycler and plots the graph (B) Detailed electrical diagram of thermocycler developed in this study.

### 2.2 Urine sample collection and isolation of bacterial plasmid

A total of 54 urine samples were collected from Meridian Health Care Center (Maharajgunj, Kathmandu) and Abhiyan Pathology Lab (Sitapaila, Kathmandu) by the hospital staffs for diagnosis purpose from patients visiting the hospitals with urinary tract infection problems. Ethical approval was approved for the use of human urine samples in this study from Nepal Research Health Council (Ref No. 688). We did not obtain personal and demographic data of patients. Samples were stored in a refrigerator at -20 ºC until used.

Isolation of plasmid DNA was carried out using a boiling method[22]. In brief, a single colony of *E. coli* isolates were taken and cultured in 5 mL LB media (Thermofisher Scientific, USA) and grown overnight at 37 ºC at 200 rpm. Overnight grown cultures were transferred to a 1.5 mL microcentrifuge tube and centrifuged at 10,000 rpm for 8 minutes[23]. Supernatant was removed and 50 μL of Tris-EDTA (TE) buffer was added to the pellet. Centrifuge tubes were then vortexed until the pellets completely dissolved. Tubes were then kept in a water bath maintained at 100 ºC for 10 minutes. Afterwards, tubes were cooled on ice and then subjected for centrifugation at about 10000 rpm for 5 minutes. Supernatant was then transferred to another tube and stored at -20 ºC until use.

### 2.2 Antimicrobial susceptibility testing

Antibiotic susceptibility tests of different isolates were performed by Kirby-Bauer disc diffusion method on Mueller Hinton agar (MHA) plates (Hi-Media Labs, India) [24]. The antibiotic discs (Hi-Media Labs, India) used for the susceptibility tests included amoxicillin (20 mcg), amoxycillin/clavulanic acid (20/10 mcg), aztreonam (30 mcg), cefotaxime (30 mcg), ceftazidime (30 mcg), ceftriaxone (30 mcg), cotrimoxazole (25 mcg), gentamicin (10 mcg), ofloxacin (5 mcg), amikacin (30 mcg), piperacillin/tazobactam (100/10 mcg) and nitrofurantoin (300 mcg). These antibiotics were chosen following standard CLSI guidelines[25]. *E. coli* isolates were regarded as multidrug resistant (MDR) if they were resistant to at least one agent of three different classes of antibiotics.

### 2.2 Combined disk test for phenotypic detection of ESBL

A combined disc diffusion test was performed for ESBL confirmation [26]. Cefotaxime (30 μg) and ceftazidime (30 μg) discs were used standalone, and with their respective combination with clavulanic acid (10 μg) for the confirmation. The MHA plates were inoculated with bacterial suspension of 0.5 McFarland turbidity standards and incubated overnight at 37 °C with both discs placed 25 to 30 mm apart [3]. An increase of ≥ 5 mm in the diameter of zone of inhibition of the respective cephalosporin/clavulanic acid combination discs than the cephalosporin discs standalone confirmed ESBL production[27].

### 2.2 Preparation and optimization of PCR conditions

DNA was extracted from urine samples using boiling method [4, 5]. Forward primer (GAGACAATAACCCTGGTAAAT) and reverse primer (AGAAGTAAGTTGGCAGCAGTG) were used for the detection of bla-TEM gene while forward primer (GAAGGTCATCAAGAAGGTGCG) and reverse primer (GCATTGCCACGCTTTTCATAG) were used for bla-CTX-M gene [6]. Similarly, we also performed reactions using conventional PCR system (BioRad T 100). Initial amplifications were carried out using the conditions from the reference [6]. However, we optimized the time condition as template size for bla-TEM and bla-CTX-M were 560 bp and 459 bp, respectively, which needed lower extension time than mentioned in the reference [7]. Table 1 shows the amplification reaction conditions for different genes in this study. PCR mixture (25 μL) consisted of 1 μL of forward primer, 1 μL of reverse primer, 10 μL of DI water, 1 μL of template DNA and 12 μL of master mix. Since our custom-built thermocycler lacked lid heating, we added 15 μL of mineral oil to prevent evaporation of the sample during operation. After amplification, gel electrophoresis (Cleaver scientific namopac 300P) was used for visualization of DNA bands. Electrophoresis was performed using 1 % agarose gel in 1 × TBE buffer for 30 minutes at 60 V. Ethidium bromide (1 μg/mL) was used for staining DNA and UV-transilluminator was used for visualization. Nuclease-free water (Sigma Aldrich, Singapore) was used as negative control during PCR experiments.

**Table 1:** PCR conditions for amplification of bla-TEM, bla-CTX-M and 16S rRNA genes

| bla-TEM | bla-CTX-M | 16S rRNA |
| --- | --- | --- |

|  | Temperature<br>(°C) | Time<br>(s) | Temperature<br>(°C) | Time<br>(s) | Temperature<br>(°C) | Time<br>(s) |
| --- | --- | --- | --- | --- | --- | --- |
| Initial denaturation | 94 | 180 | 94 | 180 | 94 | 180 |
| Denaturation | 94 | 30 | 94 | 30 | 94 | 30 |
| Annealing | 55 | 30 | 60 | 30 | 54 | 30 |
| Extension | 72 | 60 | 72 | 60 | 72 | 160 |
| Final extension | 72 | 120 | 72 | 120 | 72 | 120 |

## 3. Results and discussion

### 3.1 Performance of thermocycler

Our initial experiment on the custom designed thermocycler involved 25 µL of DI water inside PCR tube followed by the addition of 15 µL of mineral oil to prevent evaporation. Then, temperature and time conditions were provided with the use of a custom developed smartphone application. For this experiment, we used the PCR conditions for the *bla-CTX-M* gene. Serial communication was used between Arduino and computer to obtain temperature data. An open-source application “CoolTerm” was used to log temperature data at different times in a laptop. Figure 3A shows the first few cycles of thermocycler for *bla-TEM, bla-CTX-M* and 16s rRNA genes whereas the first cycle is presented in Figure 3B. Ramping rates of 0.23 °C and 0.43 °C was achieved for the heating and cooling process. PID control was used for controlling the process where 125, 5 and 1 were used for proportional, integral, and derivative constant. By using above mentioned constants, overshoot was significantly reduced for our cycle condition. Usually in temperature control system, overshoot is undesirable, especially in thermocycler where high precision is required. Annealing temperature is very critical as primers bind to un-stranded DNA at this specific temperature. However, the heating and cooling time were found to increase which increase overall process time. In our experiments, set temperatures were within ± 0.5 °C showing a good thermal stability which is significantly less than literature reported steady state error of ± 2 °C [9].

**Figure 3:**
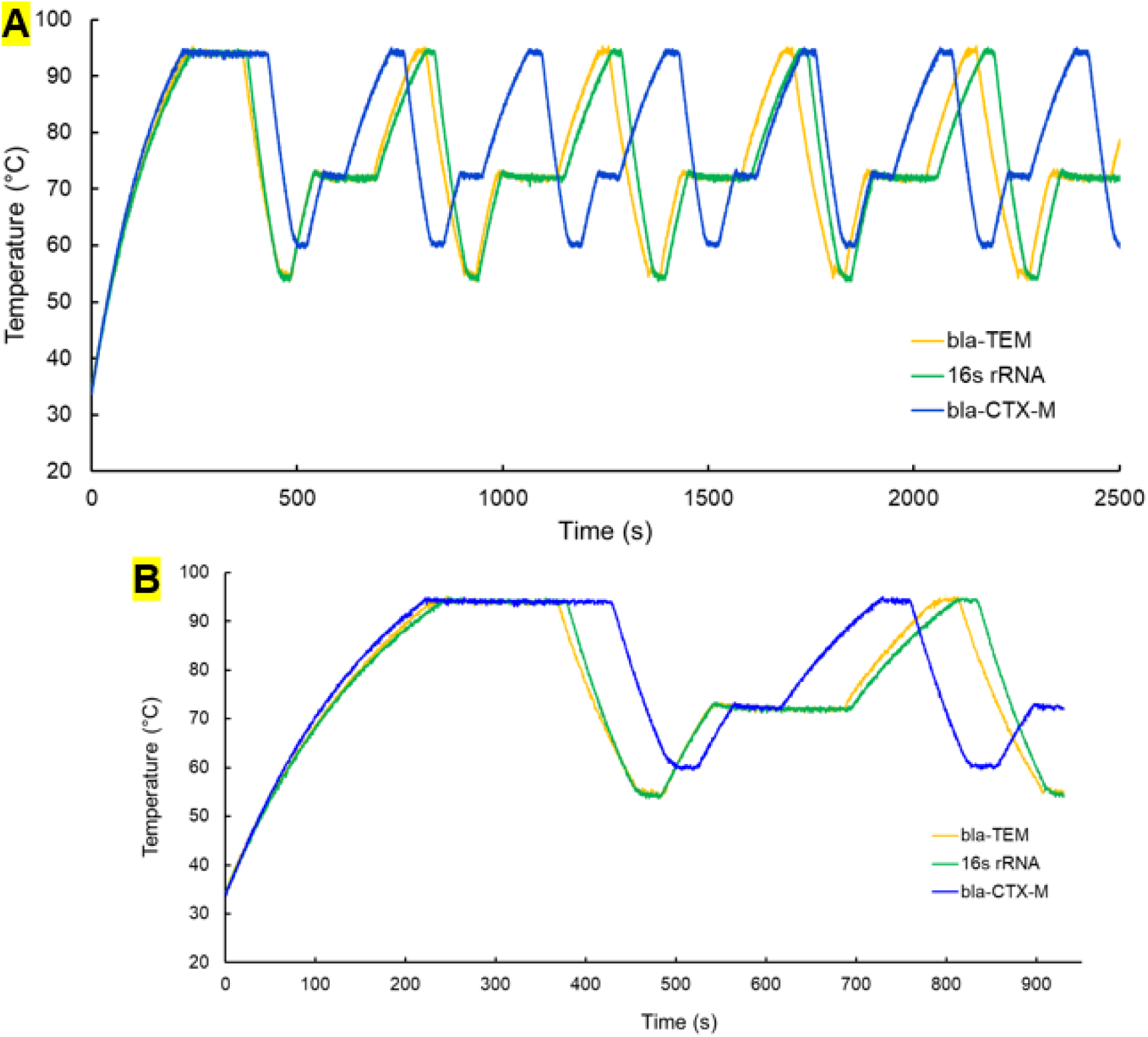
(A) Temperature cycles for bla-CTX-M, bla-TEM and 16S rRNA genes. (B) Single temperature cycle for bla-CTX-M, bla-TEM and 16S rRNA genes

### 3.2 Antibiotic susceptibility profile of *Escherichia coli*

To test the performance of custom-made thermocycler, we performed PCR of *E. coli* isolates on human urine samples. The samples were first screened for their antibiotic sensitivity. The antibiotic sensitivity test showed a high level of resistance to amoxicillin (87.1 %), amoxicillin/clavulanic acid (85.2 %), and cefotaxime (61.1 %). Bacteria were found to be highly resistant to these three antibiotics being able to hydrolyze the respective antibiotics. On either side, a high level of sensitivity was displayed in the case of amikacin (90.7 %), piperacillin/tazobactam (87.1 %), gentamicin (83.3 %), and nitrofurantoin (74.1 %) respectively. Moreover, out of 54 isolates tested, nearly 57 % of isolates were determined to be multidrug-resistant, meaning they were resistant to at least one antibiotic from three or more antimicrobial groups (Table S1). According to Yadab et al., nearly 95.5 % of *E. coli* isolates from urine samples obtained from National Kidney Hospital were found to be multidrug resistant [10]. Figure 4A shows representative photographs of discs antibiotic susceptibility testing and the results are presented in Figure 4B.

**Figure 4:**
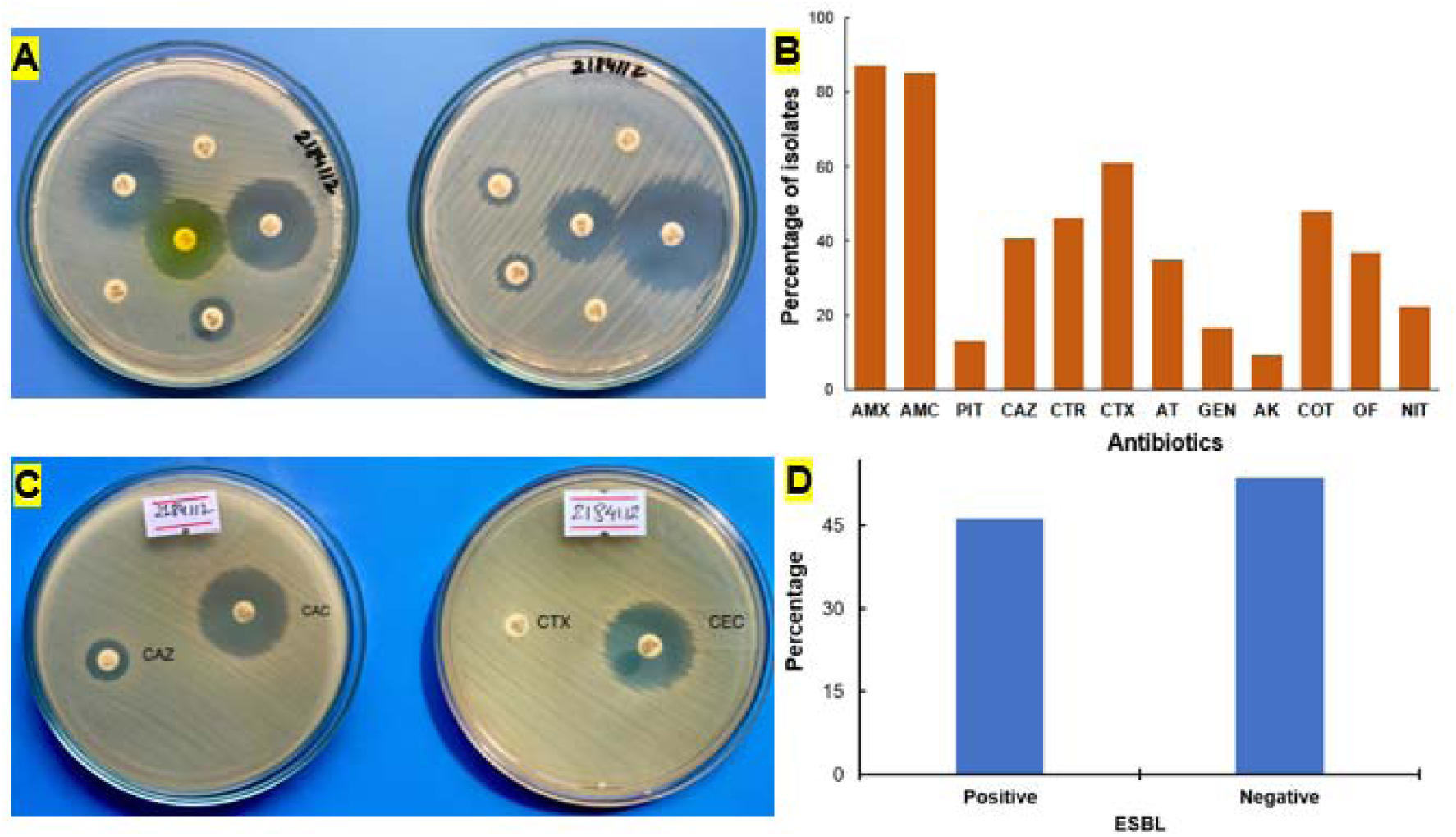
(A) Antibiotic resistance shown by *E. coli* isolate against different antibiotics. The interpretation of antibiotic resistance was done according to CLSI guidelines. (B) Resistance pattern of *E. coli* isolates to the different antibiotics. (C) Phenotypic detection of extended spectrum β-lactamase producers using cefotaxime and ceftazidime, alone and in combination with clavulanic acid (D) Screening ESBL positive according to CLSI

### 3.3 ESBL screening

ESBLs are members of the class A beta-lactamases found in gram-negative bacilli of the *Enterobacteriaceae* family which are the commonest plasmid-mediated beta-lactamases. The detection of an ESBL from a clinically significant isolate mean a therapeutic resistance to all extended spectrum cephalosporins [11]. The ESBL phenotypic confirmatory test was performed using cefotaxime (30 μg) and ceftazidime (30 μg) discs with and without clavulanic acid (10 μg). Figure 4C demonstrate ESBL screening in a petri dish whereas outcome is presented in Figure 4D. It is observed that nearly 46 % (n=54) *E. coli* isolates showed positive result for ESBL screening test whereas nearly 54 % of *E. coli* isolates tested negative. Yadav et al, found nearly 79.10 % of *E. coli* isolates to be ESBL producers from samples collected from a hospital in Nepal [10].

### 3.4 PCR amplification

All 25 presumptive ESBL-producing isolates were subjected to conventional PCR-based molecular analysis. At first, conventional PCR was used to test occurrence of two genes i.e., *bla-TEM* and *bla-CTX-M*. As reported in a literature study from Nepal, these two genes showed high prevalence in context of Nepal [29][10, 12]. For instance, *bla-TEM* showed the prevalence rate of 83.8% whereas *bla-CTX-M* had the prevalence rate of 66.1%. PCR analysis for our study showed the prevalence of *bla-*CTX-M (72%) was highest among the 25 ESBL producers, followed by *bla-*TEM (56%). The co-existence of both genes (*bla-TEM*, and *bla-CTX-M*) was seen in 11 (44%) of the isolates (*see* Table S2).

Randomly, six samples were chosen for each gene to run in the custom-built thermocycler for DNA amplification. For all the experiments, nuclease-free water was used as negative control. All six samples which showed presence of target genes using commercial thermocycler also showed positive results when using custom built thermocycler reported in this paper. The new thermocycler did not detect genes in the negative control samples. For the comparison, positive samples from both commercial PCR and custom-built PCR were used together during gel electrophoresis experiments which showed comparable performance (Figure 5A). To expand the applicability of our thermocycler, we also performed experiments to amplify 16s rRNA from *E. coli* bacteria. The reason behind this experiment is to validate if our thermocycler can amplify genes with larger size (∼1500 bp). Because of larger size of this gene, longer extension time of 2 mins 40 seconds was employed. The 16S rRNA gene was successfully amplified using our thermocycler (Figure 5B).

**Figure 5:**
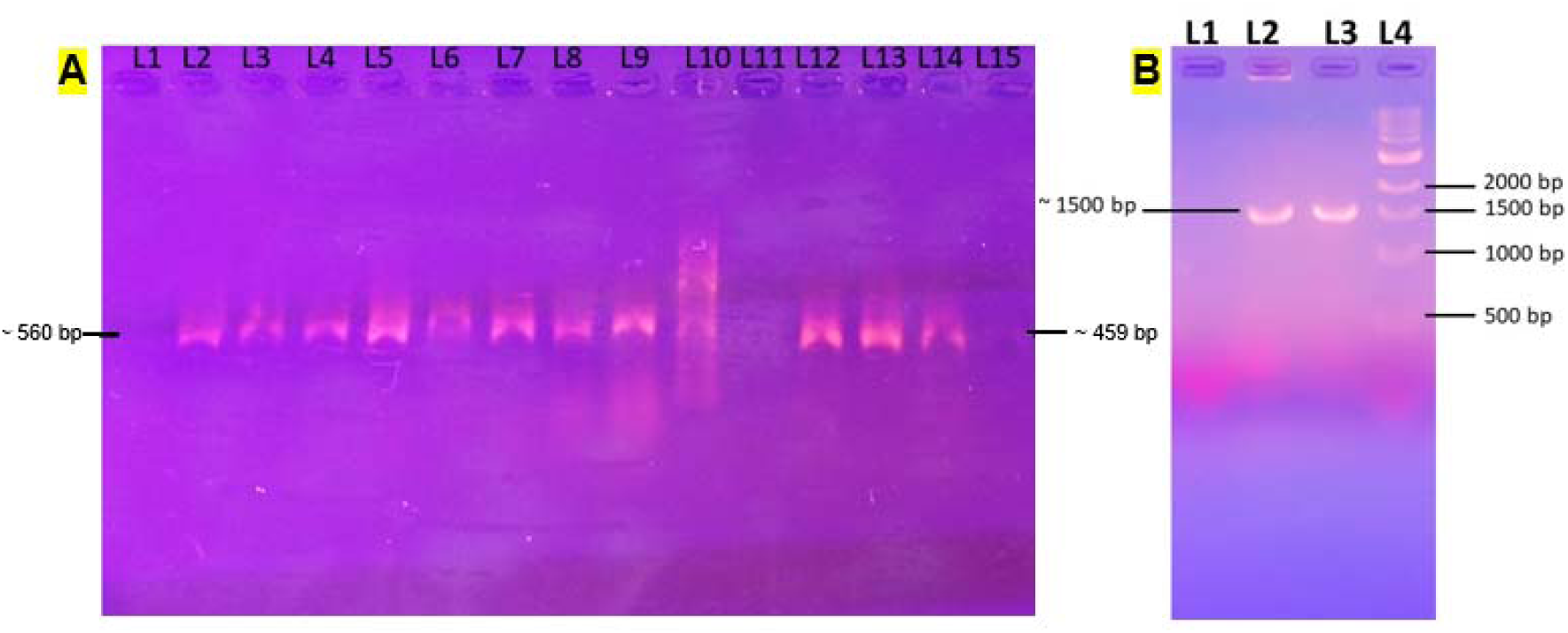
(A) Agarose gel electrophoresis (1 %) for comparing the *bla-CTX-M* and *bla-TEM* product from commercial and custom-built PCR. L1– Negative Control; L2 – L5: PCR amplicons using custom-built PCR for *bla-CTX-M* from samples 159194, 354197, 155602, 158944; L6 – L9: PCR amplicons using regular PCR for *bla-CTX-M* using the same template; L10 – 100 bp Ladder (N3231S); L11 Negative Control; L12 – L13: PCR amplicons using portable PCR for *bla-TEM* from 693125, 158871; L14 –L15: PCR amplicons using regular PCR for *bla-TEM* using the same template. (B) Gel electrophoresis result for 16s rRNA gene from custom built thermocycler

Our thermocycler was built with low cost (less than $50) which is significantly cheaper than existing commercial thermocyclers. Since this device runs on 12 V power supply, this can be operated using few 12 V batteries. Apart from this, the size of device is comparatively smaller with the size less than a kg, making this device portable, and can be used in remote location without the need of AC power supply. Moreover, temperature and time conditions can be provided wither using computer program or by using custom built smartphone application.

## 4. Conclusion

We have developed a smartphone-operated, affordable, portable, and user-friendly PCR thermal cycler. The system composed of a cartridge heater, Peltier cooling device which are controlled using Arduino Uno via the use of PID algorithm. A Bluetooth module was used to communicate with the smartphone. The reported thermal cycler can be used by anyone with minimal training. The material cost was only USD $50.0. It was operated using a wireless system using smartphone devices. There was no need of external pumps or valves. We successfully demonstrated amplification of two genes that are attributed to the production of ESBL i.e., *bla-TEM* and *bla-CTXM*. Additionally, 16s rRNA gene was also amplified using our system. It successfully detected above three genes in human urine samples with 100% accuracy. Since we only described PCR thermal cycler, it would require a gene separation and visualization system.

During COVID-19 global pandemic, we have witnessed a huge shortage and price hike of diagnostic accessories, equipment, and supplies. Affordable and user-friendly PCR thermal cycler described in this paper could contribute towards a distributed manufacturing of such systems during high demand in order to minimize the impact during emergencies. Moreover, because of low cost and lightweight (<1kg) compared to conventional PCR systems, this system can be of great use in resource limited and field setting including remote locations without stable electricity. The use of the thermal cycler can be extended, beyond the detection of AMR genes, to clinical diagnosis, genetics, forensic and environmental protection.

## Acknowledgments

We would like to acknowledge The World Academy of Sciences (TWAS), Italy (19-259 RG/CHE/AS_G to BG) for supporting M.Sc. thesis research of Garima Thapa. Experiments on commercial PCR were carried out at Research Institute for Bioscience and Biotechnology (RIBB). We thank Mitesh Shrestha from RIBB for his help.

## Ethical approval

Ethical approval was obtained from Nepal Research Health Council (Ref No. 688).

