## Supplementary information for "Smartphone-Operated Affordable PCR Thermal Cycler for the Detection of Antimicrobial Resistant Bacterial Genes"

### Smartphone-controlled low-cost thermocycler for the detection of extended spectrum beta-lactamases producing bacterial genes

#### Supplementary information

Sanam Pudasaini[1] Garima Thapa[1, 2], Bishnu P. Marasini[2], Basant Giri[1]

[1]Center for Analytical Sciences, Kathmandu Institute of Applied Sciences, Kathmandu, Nepal

[2]Department of Biotechnology, National College, Tribhuvan University, Naya Bazar, Kathmandu, Nepal

**Correspondence:**

Basant Giri

Table S1: Antibiotic susceptibility test for *E. coli* isolates

| ***Antibiotic Classes*** | **Antibiotics** | **Antibiotic Concentration (µg)** | **Sensitive %** | **Intermediate %** | **Resistant %** |
|---|---|---|---|---|---|
| *β-Lactam/ β-Lactamase inhibitors* | Amoxicillin (AMX) | 10 | 12.9 | - | 87.1 |
| | Amoxycillin/ Clavulanic acid (AMC) | 20/10 | 14.8 | - | 85.2 |
| | Piperacillin/ Tazobactam (PIT) | 100/10 | 87.1 | - | 12.9 |
| *Third generation cephalosporins* | Ceftazidime (CAZ) | 30 | 55.6 | 3.7 | 40.7 |
| | Ceftriaxone (CTR) | 30 | 50.1 | 3.6 | 46.3 |
| | Cefotaxime (CTX) | 30 | 31.5 | 7.4 | 61.1 |
| *Monobactams* | Aztreonam (AT) | 30 | 61.1 | 3.7 | 35.2 |
| *Aminoglycosides* | Gentamicin (GEN) | 10 | 83.3 | - | 16.7 |
| | Amikacin (AK) | 30 | 90.7 | - | 9.3 |
| *Folate inhibitor* | Cotrimoxazole (COT) | 25 | 50.1 | 1.8 | 48.1 |
| *Fluoroquinolone* | Ofloxacin (OF) | 5 | 61.1 | 1.8 | 37.1 |
| *Quinolone* | Nitrofurantoin (NIT) | 300 | 74.1 | 3.7 | 22.2 |

Table S2: Prevalence of *bla*$_{CTX-M}$ and *bla*$_{TEM}$ genes.

| S. N | SAMPLE CODE | ***BLA***$_{\mathbf{CTX-M}}$ | ***BLA***$_{\mathbf{TEM}}$ |
|---|---|---|---|
| 1. | 159388 | - | - |
| 2. | 155602 | + | + |
| 3. | 4991 | + | - |
| 4. | 1555 | + | + |
| 5. | 158944 | + | + |
| 6. | 156550 | - | - |
| 7. | 15546 | + | - |
| 8. | 158871 | + | + |
| 9. | N100 | + | - |
| 10. | 158564 | + | - |
| 11. | 1512123 | - | - |
| 12. | 2184112 | + | + |
| 13. | 693/25 | + | + |
| 14. | N107 | + | + |
| 15. | 159194 | + | - |
| 16. | 354197 | + | - |
| 17. | 154106 | + | + |
| 18. | 349756 | - | + |
| 19. | 155624 | + | - |
| 20. | 161572 | - | + |
| 21. | N109 | - | + |
| 22. | 1406/LF | + | + |
| 23. | 160938 | - | - |
| 24. | N110 | + | + |
| 25. | 158853 | + | + |
| TOTAL | | 18 | 14 |
